# A Preliminary list of Birds from Laguna de Matusagarati: A Wildlife Refuge in Darien Province, Panama

**DOI:** 10.1101/2024.03.14.585043

**Authors:** Jorge L. Garzón, Gumercindo Pimentel-Peralta, Pedro L. Castillo-Caballero, Jorge L. Medina, Yennifer Alfaro, Jean Paul Carrera, Anayansi Valderrama

## Abstract

The wetland Laguna de Matusagarati, in the province of Darien, eastern Panama region, possesses an approximate extension of 56, 000 hectares, the largest wetland in Panama and one of the biggest of Central America. Despite 70% of this wetland is on protected areas by Ministerio de Ambiente Panama, there are areas where the anthropogenic impact is notable, livestock, rice and oil palm crops, logging and deforestation are the most intense activities that have led to habitat loss. As part of a comprehensive One Health initiative focused on the surveillance of emerging diseases in the Darien region, we have undertaken simultaneous investigation on the biodiversity (mosquitos, birds, mammals, reptiles and amphibians) of Matusagarati. Here we report the results of the avifauna diversity undertaken in October 2018 and June 2019. We used mist nests, direct observation with binoculars and photographic cameras to report the species. A total of 169 species were reported, 49 families and 19 orders that represent 17% of the birds from Panama. Here we present the first checklist of birds from the Humedal Laguna de Matusagarati, with this we expect to improve the knowledge of the biodiversity of this wetland and provide data to authorities to improve the policies of conservation in this unique ecosystem.

---

Darien belongs to one of the 25 global mega-diversity hotspots (the Darien-Choco - Western Ecuador hotspot: Myers *et al*. 2000, Parker *et al*. 2004). This region has played a significant role in the biogeography and diversification of the neotropical region, acting as a route for or barrier to the interchange between North and South American faunas during the Tertiary and Pleistocene (Simpson 1950, Mayr 1964, Haffer 1970, Smith & Klicka 2010, Renjifo *et al*. 2017). The result of this interchange is evidenced by the diversity of organisms including birds with origins in both Central America and South America (Haffer 1967a, 1967b). Despite the high diversity of birds in Darien, the current knowledge of the avifauna is limited (BirdLife International 2014). Due the hard access to date the avifauna documented to this region is the result of expeditions (Wetmore 1965, 1968, 1972, Wetmore *et al*. 1984, Robbins *et al*. 1985, Angehr & Christian 2000, Angehr *et al*. 2004, Miller *et al*. 2011, 2014, Hruska *et al*. 2017) and the avifauna has not been evaluated properly by ornithologist (Miller *et al*. 2014).

The knowledge of the diversity of birds of Darien and the processes that create and maintain it may improve our ability to address problems in agriculture, emerging tropical diseases and conservation (Yasue *et al*. 2006, Miller 2014). Darien contains all or part of seven globally Important Bird Areas (IBA) (Angehr & Miró 2009). One of these IBAs is the Laguna de Matusagarati, the largest wetland of Panama and one of the largest in Central America covering an approximate area of 56,000 hectares which 70% is under legal protection: Filo del Tallo-Canglon hydrological reserve, the Chepigana Forest Reserve and the Humedal Laguna de Matusagarati (Carol 2020). Despite being in the managements of protected areas this wetland continues under threat due to habitat loss and fragmentation resulting from anthropogenic activities such as cattle ranching, rice and oil palm crops, and illegal hunting (CREHO 2015, Ministerio de Ambiente 2016, Jimenez 2020, Carol 2020). These activities threaten the local avifauna specially to the freshwater birds that use this wetland as breeding site (Angehr 2003). Therefore, expanding the knowledge of the avifauna of this area may improve strategies of management and conservation of this wildlife refuge.

To date the knowledge of the avifauna from the Laguna de Matusagarati is limited to a short list of birds presented in the Inventario de Los Humedales Continentales y Costeros de Panama (CREHO 2009). As part of a comprehensive One Health initiative focused on the surveillance of emerging diseases in the Darien region, we have undertaken simultaneous investigation on the biodiversity (mosquitos, birds, mammals, reptiles and amphibians) of Matusagarati. Here we present a complete checklist of the birds from the wetland Laguna de Matusagarati and information of local range extension of several species.

## Methods

### Ethical considerations

Animal research in Panama was conducted in compliance with regulatory protocols. Authorization was granted by the Panamanian Ministry of Environment under protocol SC/A-21-17, dating from February 2017. Furthermore, the Institutional Animal Care and Use Committee (IACUC) at the Gorgas Memorial Institute of Health Studies oversaw the research, operating under protocol 010/CIUCAL/ICGES18 from 2018, which adhered to the provisions of law No. 23 of January 15, 1997, governing animal welfare in the Republic of Panama.

### Study area

We surveyed two sites in the wetland Laguna de Matusagarati, Estanques de Aruza (8°21’ 53.4’’, 77°58’ 06.6’’) and Finca La Laguna (8°20’ 56.4”, 77°57’ 04.4”) (Figure 1). The wetland Laguna de Matusagarati is located near the communities of Aguas Calientes and Yaviza in Darien Province (Ministerio de Ambiente 2017). It is conformed of river margins and adjacent floodplains, includes different types of vegetation such as swamp forest, scrubland and herbaceous formation (Ibáñez & Flores 2020), a mean temperature between 21 to 25°C and an annual rainfall of 2 500 mm/year (Angehr 2003, Carrera *et al*. 2015, Ministerio de Ambiente 2016). The rainy season in this area lasts for a period of 8–9 months (May– December) and the dry period usually occurs for 3–4 months (January–April) (Carrera *et al*. 2015, Santos *et al*. 2020).

**Figure 1.**
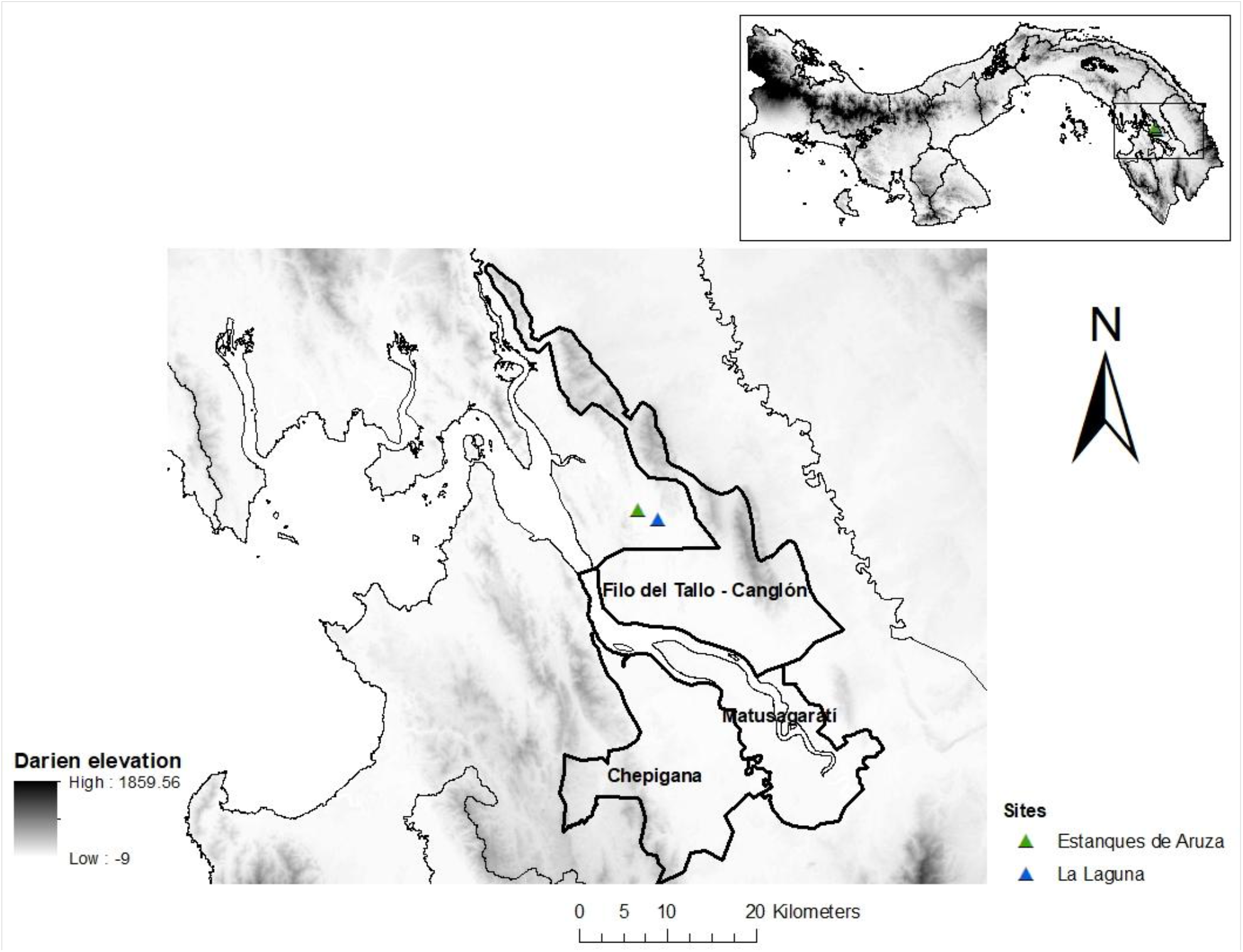
Sites sampled in the wetland Laguna de Matusagarati, Darien Province, Panama.

### Data collection

The study is part of a comprehensive One-Health surveillance initiative to detect emerging diseases in the Darien region. We have undertaken simultaneous investigation on the biodiversity (mosquitos, birds, mammals, reptiles and amphibians) of Matusagarati. Birds were caught in two periods, the first from 15 – 17 October 2018 at the site Finca La Laguna and the second from June 29 to July 1, 2019, at Estanques de Aruza. We spent 3 days at each site using 12 × 5 m nylon mist nets open from 7:00 to 17:00 hours depending on the bird activity. Once trapped, birds were extracted into paper bags and then transferred into fabric bags to avoid ectoparasite crossover. We also recorded birds using binoculars and cameras during our walk to and from the study sites from 6:00 – 18:00. We used the Bird of Panama A Field Guide (Angehr & Dean 2010) to identify the species in the field and evaluate the data of range extension. The American Ornithologist Union checklist was followed to ensure updated scientific names (Chesser *et al*. 2023).

## Results

A total of 169 bird species distributed across 49 families and 19 orders were reported, representing 17% of the recognized species in Panama (AUDUBON 2023). The most numerous orders were Passeriformes with 19 families and 81 species. The number of species reported through mist nets were 49, while the other 120 species were reported using binoculars and cameras. During the migratory season in October 2018, 14 migratory species were reported. According to the IUCN Red List 2020, one species is under the Vulnerable (VU) category and another Near Threatened (NT). The List of threatened species of flora and fauna from Ministerio de Ambiente Panamá (2016) listed 21 of the species observed as VU and one as Critically Endangered (CR).

Here we provide a list of species with local range extension based in the Bird of Panama A Field Guide (Angehr & Dean 2010).

### BLACK-CROWNED NIGHT-HERON (*Nycticorax nycticorax* Linnaeus, 1758)

We observed one individual in June 2019 resting among the dense vegetation on the lagoon next to the large colony of Cattle Egrets (fig. 2B). This species nests individually or in mixed species colonies of other herons and ibises (Kushlan & Hancock 2005). The Black-crowned Night-Heron has often been used as an indicator of environmental good quality because it is an upper trophic level bird that nests colonially, has a wide geographic distribution, and tends to accumulate contaminants (Custer *et al*. 1991). This species was reported as part of the fauna in the Laguna de Matusagarati in the Inventario de los Humedales de Panamá (CREHO 2009). Wetmore *et al*. (1965) also reported the Black-crowned Night-heron in Darien, but this species shares the common name “hurafia” with Yellow-crowned Night-heron, and it is unclear whether both were seen. The distribution of Black-crowned Night-heron described in the Birds of Panama A Field Guide by Angehr & Dean (2010) does not include this region of the Darien Province.

**Figure 2.**
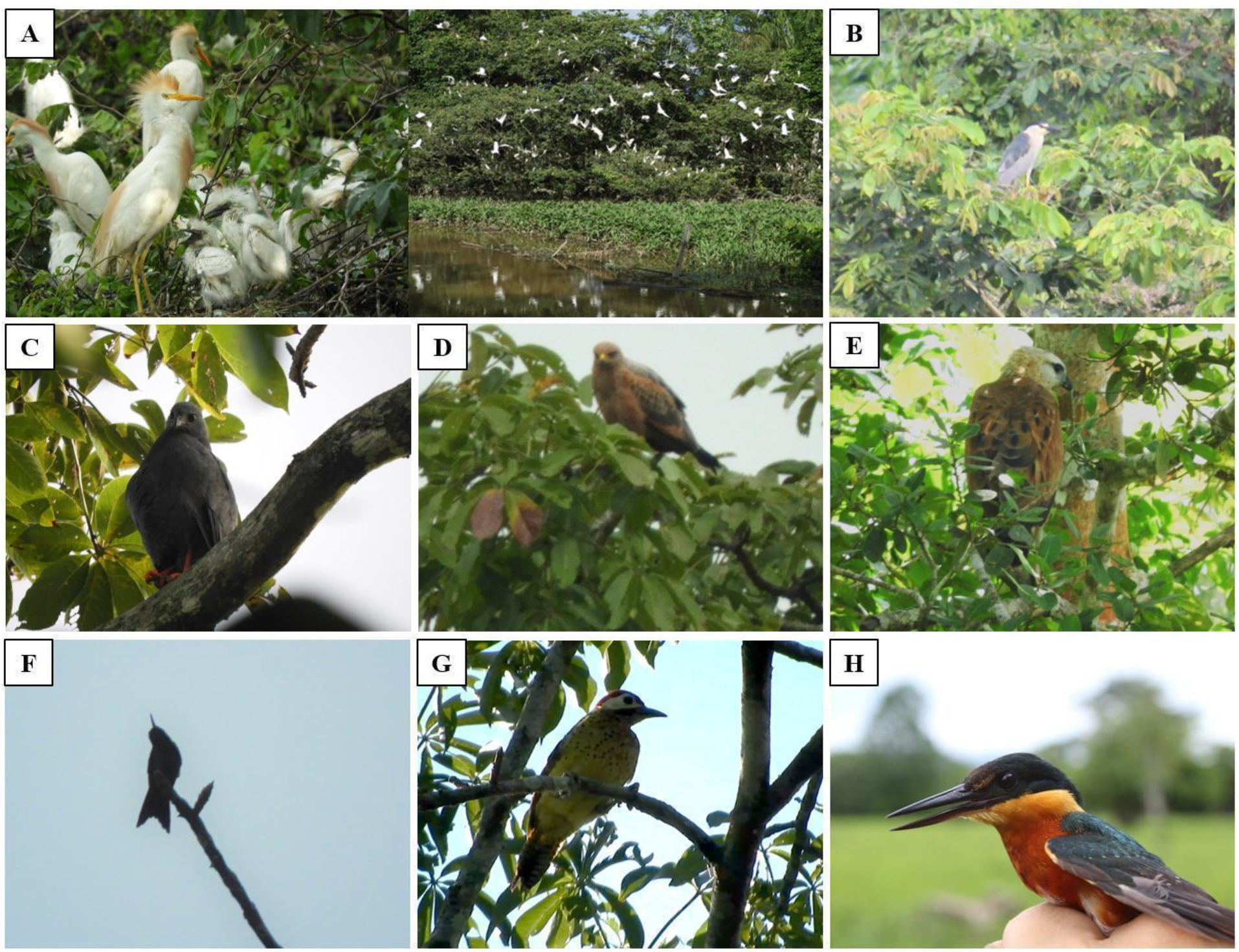
Important bird species observed in the Wetland Laguna de Matusagarati in 2018-2019; colony of *B. ibis* nesting (A), *N. nycticorax* (B), *G. caerulescens* (C), *B. meridionalis* (D), *B. nigricollis* (E), *T. colombica* (F), *C. punctigula* (G), *C. aenea* (F).

### SNAIL KITE (*Rostrhamus sociabilis* Vieillot, 1817)

We saw one individual near our mist netting site in October 2018. This report is the first for the wetland Laguna de Matusagarati and second for Darien Province, this last was reported in Río Marea by Chris Fisher on November 23, 2017 (Fisher & Arias 2017, eBird checklist: S40682994). The first report for Panama was on March 22, 1929, in Permé by Wedel (Wetmore *et al*. 1965). Angehr & Dean (2010) report the distribution of this species as restricted to the canal zone.

### SAVANNA HAWK (*Buteogallus meridionalis* Latham, 1790)

In 2018 and 2019, we observed several individuals walking, perching, and flying over the areas near a pond (fig.2D). No records were previously known from Darien (Ridgely & Gwynne 1989, 1993). According to Angehr & Dean (2010), this medium-sized raptor’s distribution comprises most of Panama’s Pacific slope, extending to a small part to the north-west of the Darien Province. It has only recently (since 2008) spread to SW Costa Rica as a result of ongoing deforestation (Sandoval *et al*. 2010). This account is part of an increase in reports of this species in the Darien lowlands over the last 10 years (eBird 2021).

### BLACK-COLLARED HAWK (*Busarellus nigricollis* Latham, 1790)

We observed an adult near the camp, perched in a tree on the edge of the Finca La Laguna in October 2018 (fig.1E). Despite its wide distribution across the Americas, Panama has a localized distribution in its eastern regions, with few historical records in the central Panama provinces of Veraguas and Coclé (Ridgely & Gwynne 1993, Angehr & Dean 2010). Panamanian populations have been declining due to drainage of wetlands, and the same may be true elsewhere (Bierregaard *et al*. 2020). Populations are only known in the east of the country (from the Canal Zone to the Darien Province) according to the distribution map of the Bird of Panama A Field Guide by Angher & Dean (2010) except for a single record on the north-west side of the country very close to Costa Rica by Decastro González in 2015 (eBird Checklist: S25348643). Wetmore *et al*. (1965) mentioned Festa collected an individual in Darien, Laguna Pita in 1895. Another individual was collected by J.L. Baer in 1924 near Yaviza, and an individual was observed upstream near to the mouth of the river Tuqueza resting in a tree in a marshy forest. This species is under the CR category in The List of threatened species of flora and fauna, Ministerio de Ambiente Panamá 2016.

### CROWNED WOODNYMPH (*Thalurania colombica* Bourcier, 1843)

In July 2019, we observed a male in Laguna de Matusagarati site (fig.2F). Although this species was reported by Hruska *et al*. (2016) in Darien National Park, it is not shown to occur in Darien following the Birds of Panama A Field Guide by Angehr & Dean (2010), so this report supports Hruska’s 2016 account of a species from Darien. This species is listed as Vulnerable in the List of threatened species of flora and fauna, Ministerio de Ambiente Panamá (2016).

### YELLOW-HEADED CARACARA (*Milvago chimachima* Vieillot, 1816)

We observed 3 individuals in grassy, open areas and near the edges of the pond in our walk around the Estanques de Aruza (fig.3B). Although this species is well known in Panama, we found no reports of this bird in Darien. This is the easternmost report for this species in Panama consulting The Bird of Panama a Field Guide. The first record in Darien Province was made in Río Piñas by Miller *et al*. (2011). This species previously had only been reported as far east as the Panama Province (Río Bayano) by Wetmore in 1965, Ridgely & Gwyanne 1989, 1993, Angehr & Dean 2010.

**Figure 3.**
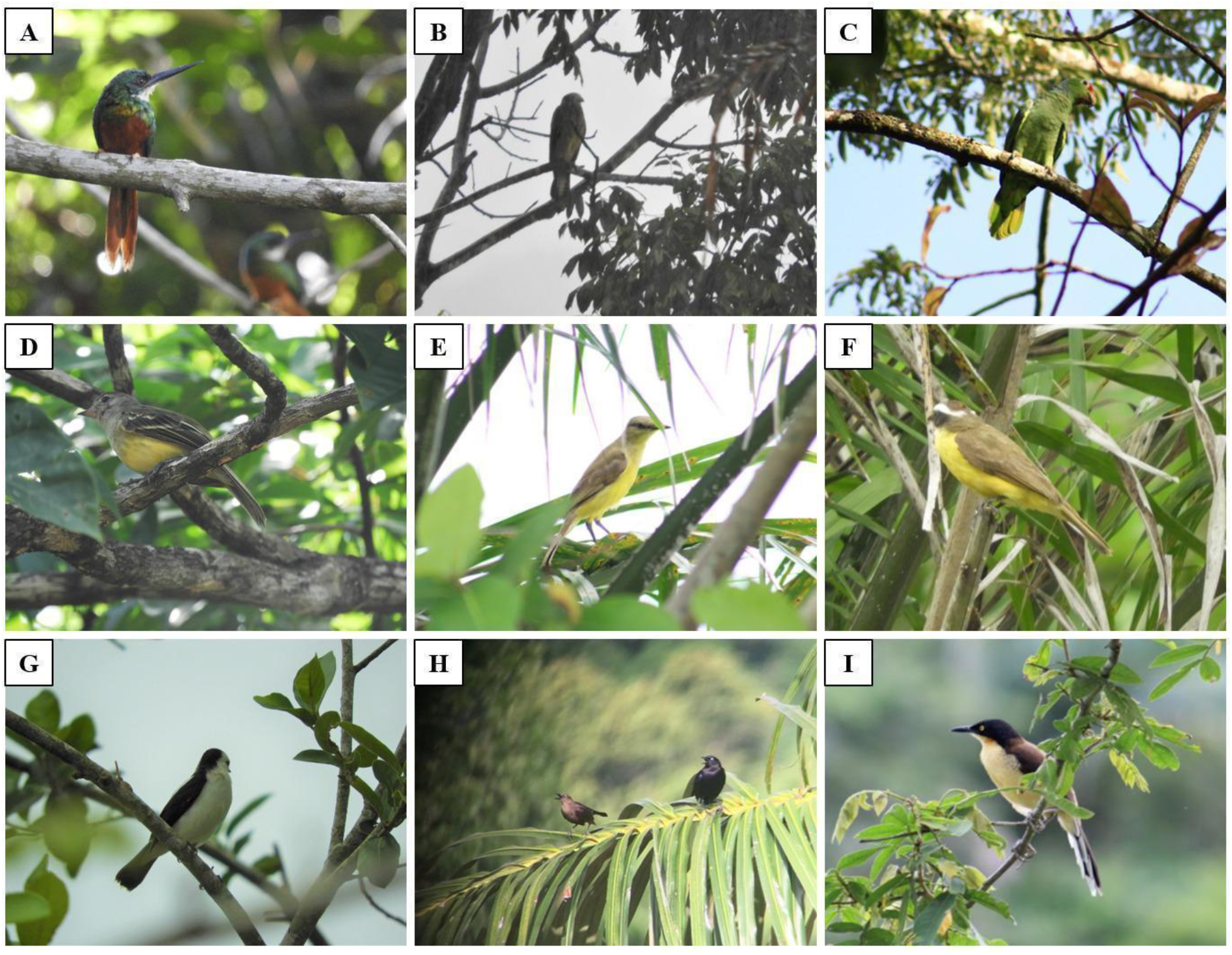
Important bird species observed in the Wetland Laguna de Matusagarati in 2018-2019.; *G. ruficauda* (A), *M. chimachima* (B), *A. autumnalis* (C), *E. chiriquensis* (D), *M. rixosa* (E), *M. similis* (F), *F. pica* (G), *Q. lugubris* (H), *D. auricapilla* (I).

### LESSER ELAENIA (*Elaenia chiriquensis* Lawrence, 1865)

We heard an individual vocalizing in some bushes in July 2019 (fig. 2D). There are only 3 records in Darien Province; the first report was in Punta Patiño, Darien by Beck & Borgmann (2012) eBird checklist: S13866414. It is frequently found in areas with grasslands and small shrubs in the lowlands of the Pacific slope from Chiriqui to the eastern regions of Panama Province (Ridgely & Gwynne 1993, Angehr & Dean 2010).

### SOCIAL FLYCATCHER (*Myiozetetes similis* Spix, 1825)

We photographed an individual on July 4, 2019, while it perched on emergent vegetation in a pond at noon (fig.3F). The distribution of this flycatcher in Panama extends to the eastern part of the Darien Province on the Pacific side (Angehr & Dean 2010). Wetmore (1972) found no evidence for this species in Darien and Guna Yala, just an observation by D. Sheets near the Panama-Colombia border in Puerto Obaldia and La Bonga.

### CARIB GRACKLE (*Quiscalus lugubris* Swainson, 1838)

This species is a recent arrival to Panama. We report four individuals in 2019 during our walk around Estanques de Aruza (fig.3H). The first report for Panama was made by Cubilla on August 15, 2017, in Finca Bayano, Darien (eBird Checklist: S38658238). Carib grackles are omnivorous birds that rest in communal groups in the middle part of the canopy and are often seen nesting in the middle parts of trees and palms (Ayerbe-Quiñones 2018).

### BLACK-CAPPED DONACOBIUS (*Donacobius atricapilla* Linnaeus, 1766)

During our walk around the Estanques de Aruza site, we photographed a pair of this species (fig.2I). This species originates from South America and the eastern Darien province, this report is the westernmost for the species in Panama. The Black-capped Donacobius is a familiar bird often seen in marshes and wet pastures across much of South America, calling attention to itself with loud, duetting calls. The subspecies *brachypterus* occurs in Panama (eastern Darien) and northwest Colombia (Kroodsma & Brewer 2020). It was reported in Serranía de Pirre, Darien Province by Robbins *et al*. (1985).

## Discussion

### Biodiversity

The checklist of birds here highlights the importance of the wetland Laguna de Matusagarati as a site of high diversity, with 17% of the birds of Panama. We present a complete list of birds reported in this wetland, some species with local range extension such as *R. sociabilis, Geranospiza caerulescens, B. meridionalis, M. chimachima, Chloroceryle aenea, Galbula ruficauda, E. chiriquensis, M. similis, Fluvicola pica, Donacobius atricapillus*; and several listed under a threat category in both national and international organizations (Ministerio de Ambiente Panamá 2016b, IUCN Red List 2020). The recent observation of the globally “Near Threatened” Northern Screamer by Pizarro in 2020 (eBird checklist: S69033434) just a few kilometers from the Laguna de Matusagarati encourages to continue monitoring this area to report birds that may be arriving from South America and can established for example the species *D. atricapilla*.

### Conservation

The wetland Laguna de Matusagarati is important for birds, providing habitat for breeding, nesting, rearing young birds, feeding, resting, shelter and social interactions. The results of this survey improve the knowledge of the avifauna of this wetland and evoke to national and international authorities the need to increase the conservation measures due to the high numbers of species reported under certain categories of threat. We recommend environmental organizations and conservationists continue monitoring the fauna and flora to provide data that can be used to improve the policies of conservation for this important wetland. Finally, this checklist may aid in developing birding ecotourism and help the communities that depend on this wetland.

## Acknowledgments

We are very thankful to the STRI Bird Collection, who provided material to complete field work. This work was supported by SENACYT, through the grants number FID-16-201 and FID-2021-96 grant to JPC; JLG was supported by the SENACYT scholarship program BIDP-2018-2019.

---

**Table 1.**
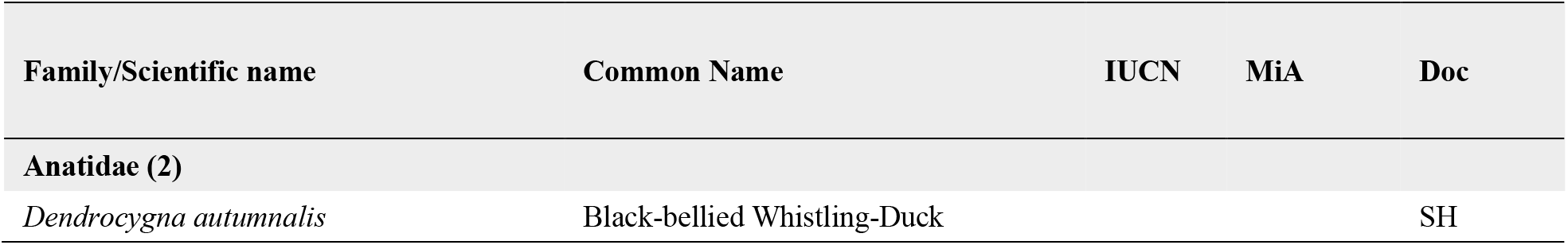
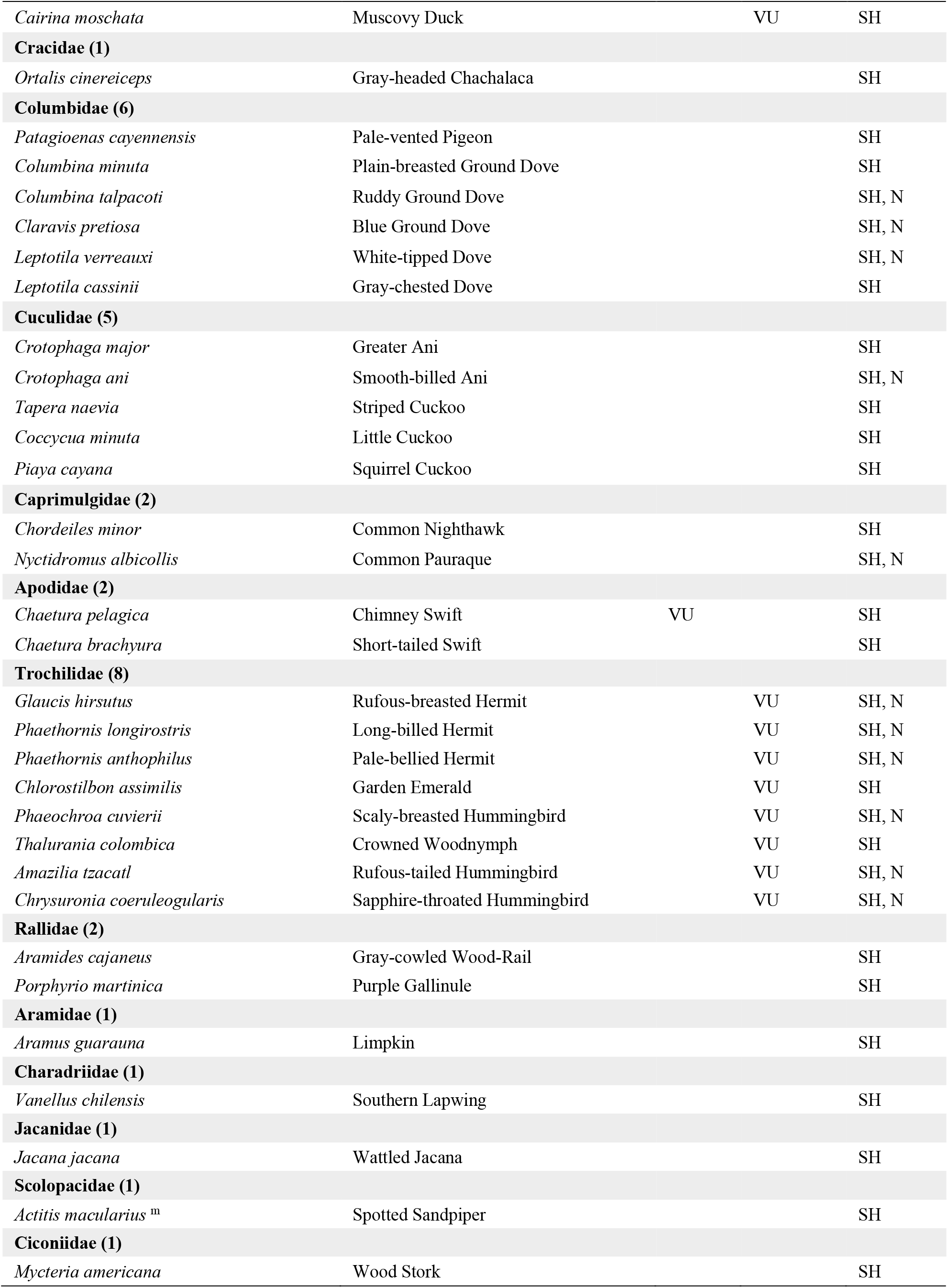
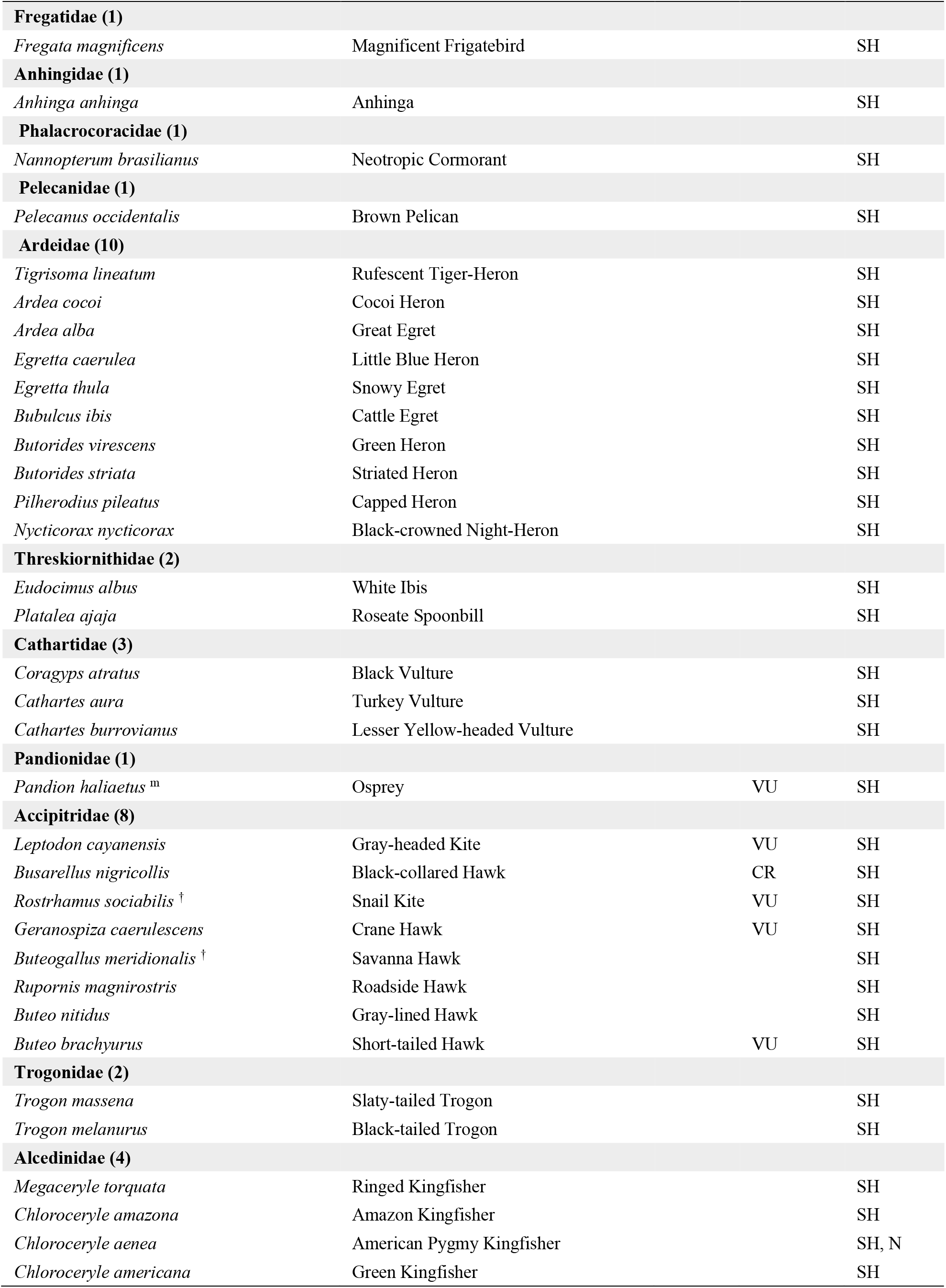
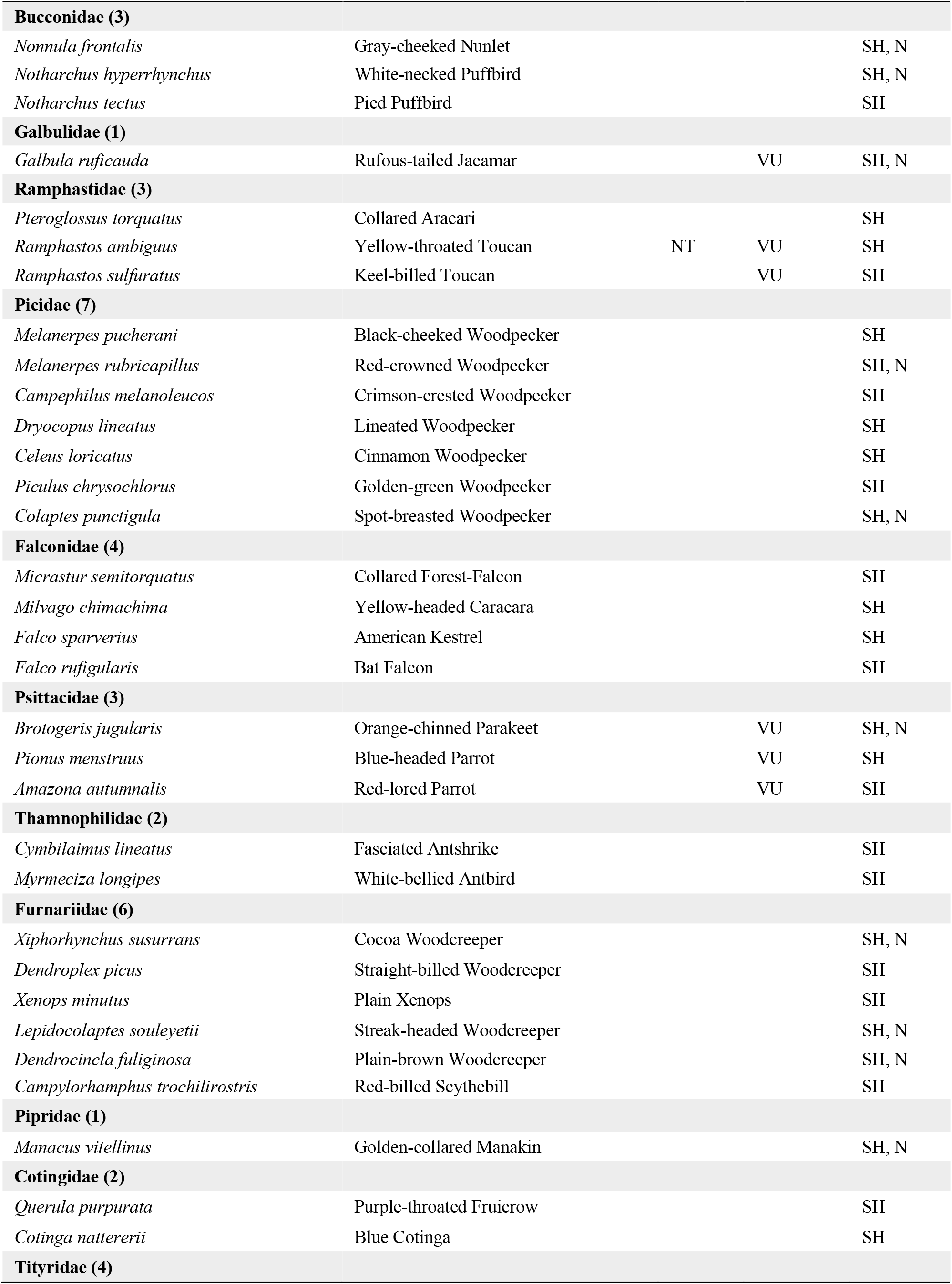
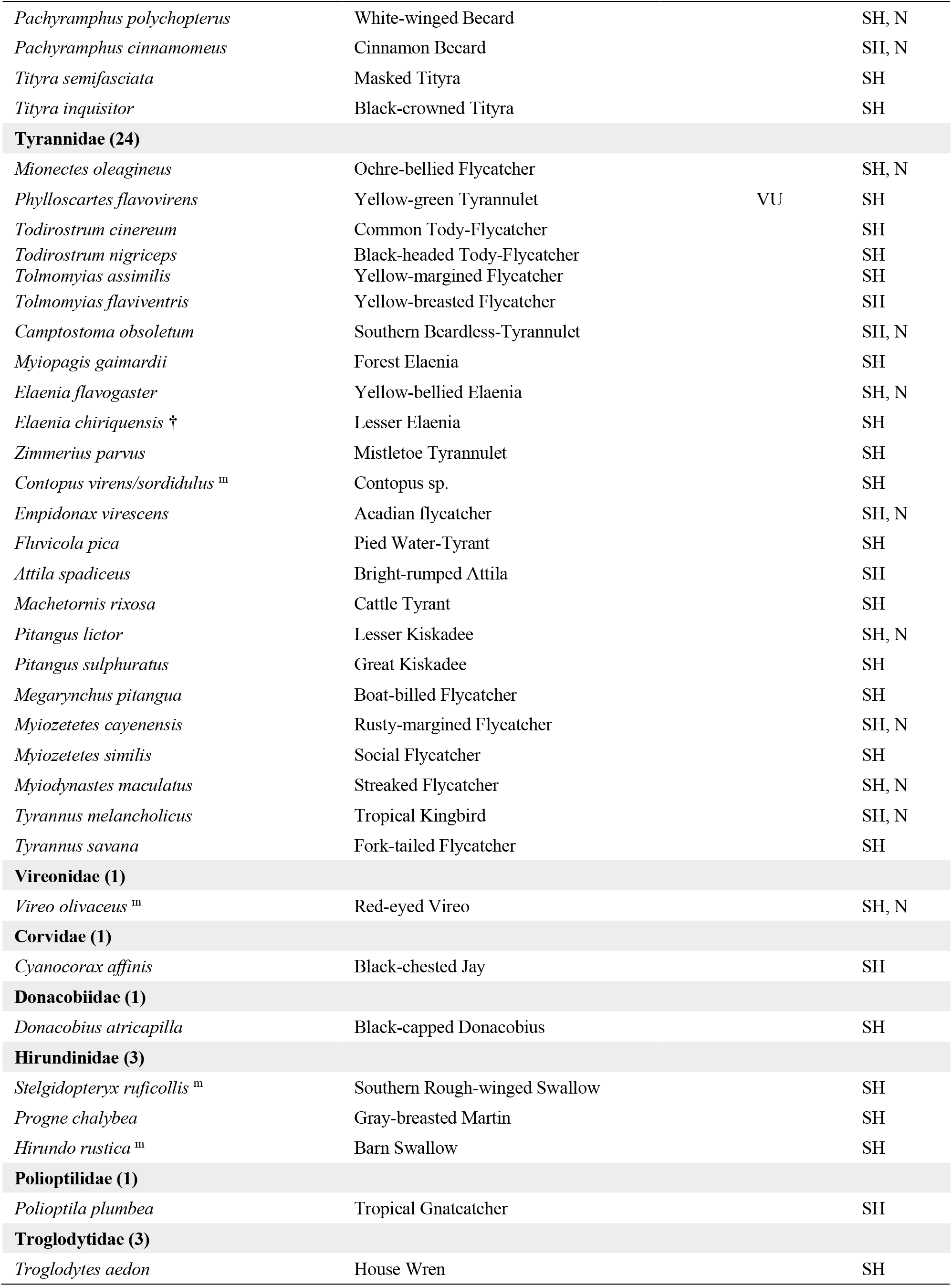
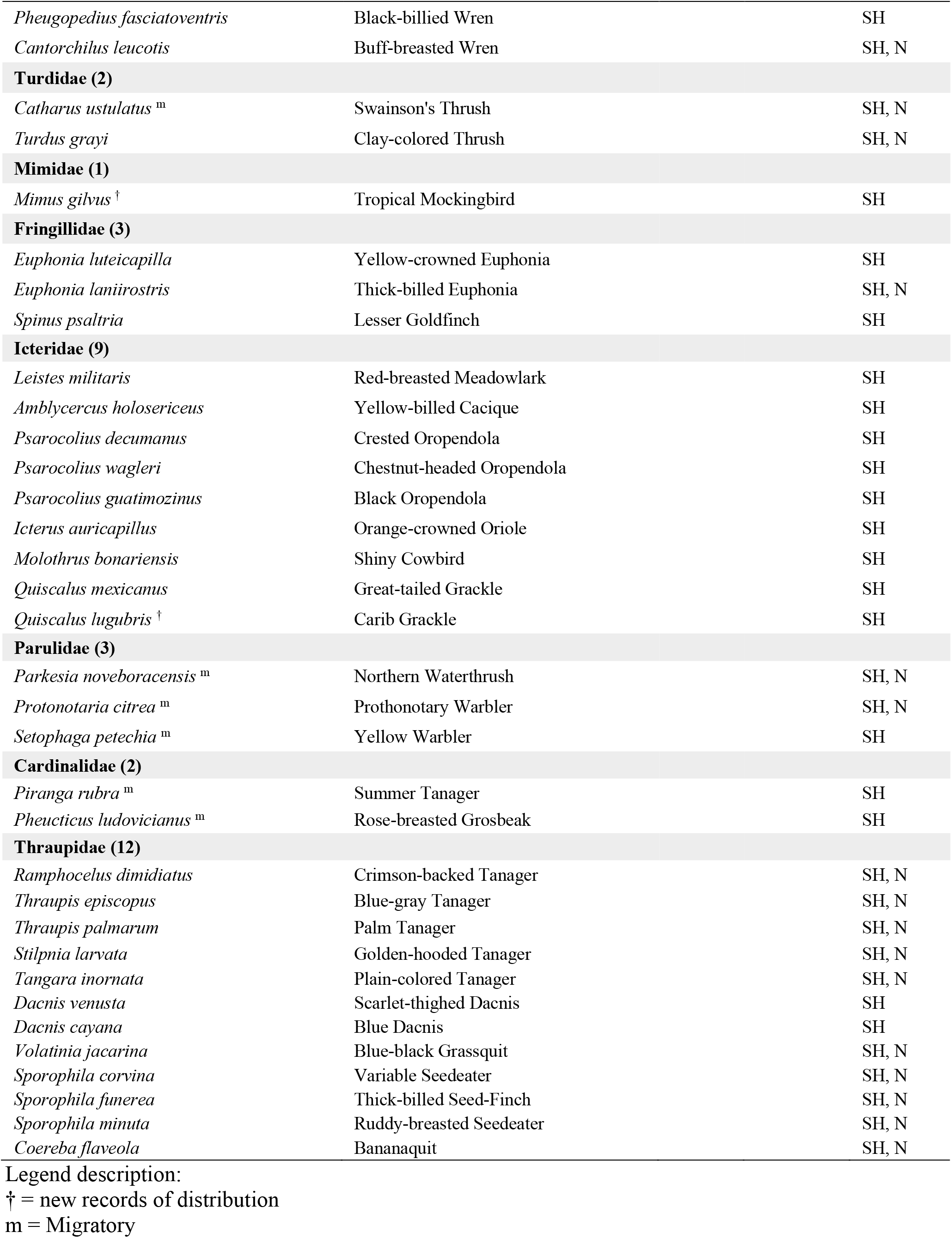
List of the avifauna community at Matusagarati ponds. IUCN and MiAmbiente (MiA) Status: LC – Least Concern (left as blank spaces); NT – Near Threatened; VU – Vulnerable. The number of species per family are provided in brackets; Documentation: SH = seen or heard only; N = caught in nets.

